# *HAC1* contributes to stress adaptation and virulence in the emerging fungal pathogen *Candida auris*

**DOI:** 10.64898/2026.03.19.712804

**Authors:** Sayoko Oiki, Masahiro Abe, Ai Hirasawa, Ami Koizumi, Amato Otani, Takayuki Shinohara, Yoshitsugu Miyazaki

## Abstract

*Candida auris* (*Candidozyma auris*) is an emerging multidrug-resistant fungal pathogen that poses a significant global health threat. However, the molecular mechanisms underlying its virulence remain incompletely understood. In this study, we performed *in vivo* transcriptome analysis using an immunosuppressed mouse gastrointestinal infection model to identify genes associated with host-adaptation and virulence during infection. By comparing fungal transcriptomes obtained from colonization and dissemination sites with those from *in vitro* cultures, we identified genes that were consistently upregulated during infection. Among these genes, the unfolded protein response regulator *HAC1* was selected as a candidate virulence-associated gene for further analysis. RT-PCR and sequencing analyses revealed that *HAC1* mRNA in *C. auris* undergoes an unconventional splicing event of 287 bp that is enhanced under ER stress conditions. The excised region spans the annotated open reading frame boundary, suggesting that the translated region of *HAC1* may require re-evaluation. Notably, a proportion of *HAC1* transcripts appeared to be spliced even under non-stress conditions, indicating a detectable basal level of UPR activation. Differences in splicing dynamics were also observed among clade strains. Functional analyses demonstrated that deletion of *HAC1* increased sensitivity to ER stress and heat stress. The *HAC1* deletion mutant also exhibited reduced virulence in both *Galleria mellonella* and immunosuppressed mouse infection models, as evidenced by delayed host mortality and decreased fungal burdens, respectively. These findings indicate that *HAC1* contributes to ER stress adaptation, thermotolerance, and survival in the host environment, and identify *HAC1* as a virulence-associated gene in *C. auris*.

## Introduction

Invasive candidiasis represents a major threat to human health, particularly among immunocompromised individuals, leading to high mortality and morbidity worldwide^1,2^. Among pathogenic *Candida* species, *Candida auris* (*Candidozyma auris*) has recently emerged as a significant global health concern due to its multidrug resistance, ability to cause nosocomial outbreaks, and persistence in healthcare environments. Since its first identification in 2009^3^, *C. auris* has been reported worldwide and is now recognized as a global public health threat^4^. The Centers for Disease Control and Prevention (CDC) and the World Health Organization (WHO) have designated *C. auris* as a priority fungal pathogen requiring urgent attention^5,6^. Genomic studies have revealed that *C. auris* is divided into five clades with different geographic distributions. These clades include clade I (South Asia), clade II (East Asia), clade III (Africa), clade IV (South America), and clade V (reported from Iran), and exhibit phenotypic diversity, including differences in virulence and antifungal susceptibility^5,7^.

The pathogenic mechanisms of *Candida* species have been extensively studied in the model pathogen *Candida albicans*. Multiple virulence factors have been identified in this species, including adhesins (agglutinin-like sequence [ALS] proteins), morphogenetic regulators (Hwp1, Efg1, and Cph1), secreted enzymes (secreted aspartyl proteases [SAPs]), and stress response pathways (Hog1 and Cap1)^8–12^. In contrast, the virulence mechanisms of *C. auris* remain incompletely understood. Recent studies have identified several genes that may contribute to the virulence of *C. auris*, such as Als4 (promoting host adhesion and virulence) and Scf1 (promoting host colonization and surface adherence)^13,14^. However, the broader molecular basis of host adaptation and virulence in *C. auris* remains unclear.

The unfolded protein response (UPR) is a conserved cellular pathway that contributes to fungal adaptation to host environments by maintaining endoplasmic reticulum (ER) homeostasis under conditions of protein-folding stress. Accumulation of misfolded proteins in the ER triggers ER stress and activates the UPR, which induces genes that enhance ER protein-folding capacity and restore cellular homeostasis. In fungi, the transcription factor Hac1 serves as the central regulator of the UPR. Activation of Hac1 depends on unconventional splicing of *HAC1* mRNA, in which the ER-resident sensor Ire1 catalyzes a non-canonical splicing reaction in response to ER stress. This splicing event enables production of the active basic leucine zipper (bZIP) transcription factor Hac1, which induces genes involved in protein folding, ER quality control, protein secretion, and degradation of misfolded proteins^15,16^. The Hac1-dependent UPR pathway has been extensively characterized in the model yeast *Saccharomyces cerevisiae* and is broadly conserved among fungi, including *Aspergillus fumigatus*, *Cryptococcus neoformans*, and *Candida albicans,* in which it has been implicated in adaptation to stresses imposed by the host during infection^17–20^. However, the role of the Hac1-mediated UPR pathway in the emerging pathogen *C. auris* remains poorly understood.

In the present study, we performed *in vivo* transcriptome analysis to identify genes induced during infection by *C. auris* and selected Hac1, an ER stress-responsive transcription factor, for further analysis. We further investigated the splicing mechanism and stress-related functions of Hac1, and evaluated its contribution to virulence using infection models. Our findings would provide new insights into the role of the UPR pathway in the virulence of *C. auris*.

## Materials and Methods

### Strains and culture conditions

*Candida auris* AR 0390 (clade I), AR 0381 (clade II), AR 0384 (clade III), AR 0385 (clade IV), and AR 1097 (clade V) were provided by the Centers for Disease Control and Prevention AR Bank. *C. auris* cells were cultured overnight in yeast extract peptone dextrose (YPD) medium at 30°C with agitation at 1,000 rpm in an RTS-8 personal bioreactor (Biosan Ltd., Latvia).

### Host animals

Female 5-week-old C57BL/6J and BALB/c mice were obtained from Japan SLC Inc. (Shizuoka, Japan) and maintained under specific pathogen-free conditions. *Galleria mellonella* larvae were purchased from Spheroaqua Inc. (Shizuoka, Japan) and maintained at 15°C in the dark until the experiments. All animal experiments were performed using a protocol approved by the National Institute of Infectious Diseases (approval numbers: 124052 and 125149). All staff were sufficiently educated in animal care and handling before this procedure.

### RNA-seq

An intestinal colonization and dissemination mouse model was established as previously described ^21^. Briefly, C57BL/6J mice were given drinking water containing an antibiotic cocktail consisting of vancomycin (45 mg/L), gentamicin (35 mg/L), kanamycin (400 mg/L), metronidazole (215 mg/L), and colistin (850 U/L) for at least 1 week before inoculation with *C. auris*. Cortisone acetate (225 mg/kg) was administered subcutaneously on 1 day before, on the day of, and 1 day after *C. auris* inoculation. A 100-µL suspension of *C. auris* AR 0390 (1.0 ×□10^9^ cells/mL) was orally administered using a sterile feeding needle. Fourteen days post-infection, mice were euthanized and intestinal contents and kidneys were collected and immediately frozen in liquid nitrogen. Total RNA was extracted from frozen intestinal contents and kidney samples using ISOGEN (NIPPON GENE) and purified using an RNeasy Mini Kit (QIAGEN) according to the manufacturer’s instructions. RNA sequencing libraries were prepared and sequenced by Azenta (South Plainfield, NJ, USA) using an Illumina platform to generate paired-end reads. Raw sequencing reads were mapped to the *C. auris* reference genome (GCA_002759435.3). Gene expression levels were calculated as FPKM values. Downstream analyses, including principal component analysis (PCA), heatmap visualization, and identification of differentially expressed genes, were performed using the iDEP program^22^.

### Construction of mutants

Deletion mutants were constructed using an RNP-based CRISPR–Cas9 system as previously described^23^, with minor modifications. To construct a repair template, the nourseothricin resistance gene *NAT1*, flanked by 100-bp homologous sequences corresponding to the upstream and downstream regions of the *HAC1* open reading frame (ORF), was amplified by polymerase chain reaction (PCR) using the pLNMCL plasmid as a template^24^. The crRNA targeting the *HAC1* ORF was designed using CHOPCHOP and synthesized by Integrated DNA Technologies, Inc (IDT). To create the Cas9-RNP complex, crRNA and tracrRNA were mixed with nuclease-free water to a final concentration of 4 µM each and incubated at 95°C for 5 min. A 3.6 µL aliquot of the resultant gRNA was mixed with 3 µL of 4 µM Cas9 nuclease V3 (IDT) and incubated for 5 min at room temperature. To prepare competent cells for electroporation, *C. auris* cells were cultured in 50 mL YPD medium at 30°C until reaching an optical density at 600 nm (OD_600_) of 1.6–2.2. Cells were harvested by centrifugation at 2,000 × *g* for 5 min, resuspended in 10 mL of transformation buffer (100 mM lithium acetate, 10 mM tris(hydroxymethyl)aminomethane (Tris-HCl), and 1 mM EDTA), and incubated at 30°C and 180 rpm for 1 h. After addition of 250 µL of 1 M dithiothreitol (DTT; Wako Pure Chemical Corporation), incubation was continued for 30 min under the same conditions. Cells were washed twice with cold water and once with cold 1 M sorbitol, then resuspended in 400 µL of cold 1 M sorbitol. For transformation, 50 µL of the competent cells were mixed with 1 µg of the repair template and 6.6 µL of RNP complex and transferred to a cold 0.2 cm electroporation cuvette (NEPA GENE). Electroporation was performed using an Electro Cell Manipulator 600 (BTX) at 1.8 kV, 200 Ω, and 25 µF. Immediately after pulsing, 1 mL of cold 1 M sorbitol was added, and cells were collected and resuspended in YPD medium. After the incubation at 30°C for 3 h for cell recovery, the electroporated cells were plated on YPD agar containing 100 µg/mL nourseothricin (Wako Pure Chemical Corporation) and incubated at 30°C for 2 days. The resulting transformants were screened by PCR to confirm deletion of the target gene.

### Construction of complement strain

To construct the complement strain, the same RNP-based CRISPR-Cas9 system used for generating the mutant strain was employed. As the repair template, a DNA fragment containing the ORF of the target gene together with the 100-bp upstream and 600-bp downstream regions was amplified by PCR from the wild-type genomic DNA and fused with the hygromycin resistance gene HygR, which was synthesized based on the sequence reported for the pYM70 plasmid^25^. Electroporation of competent cells with the RNP complex and repair template was performed as described above, and transformants were obtained on YPD agar containing 600 µg/mL hygromycin. Transformants exhibiting hygromycin resistance but sensitive to nourseothricin were subsequently screened, and proper complementation of the target gene was confirmed by PCR.

### Infection model using Galleria mellonella larvae

To screen for virulence, *G. mellonella* larvae were used as an infection model. A 10-µL aliquot of a *C. auris* cell suspension (1.0 ×□10^8^ cells/mL) prepared from the wild-type strain (AR 0390) or the mutants was injected into the last proleg of each larva using a microsyringe (Trajan Scientific Japan). Control larvae were injected with PBS. After injection, larvae were incubated at 37°C for 7 days, and survival was recorded daily. Ten larvae were used per group, and the experiments were performed independently three times. Virulence was analyzed using Kaplan–Meier survival analysis.

### Sensitivity to various stresses

To assess stress sensitivity, *C. auris* cells in the logarithmic growth phase were harvested from YPD medium by centrifugation, washed with PBS, and diluted tenfold to concentrations ranging from 1.0 ×□10^8^ to 1.0 ×□10^3^ cells/mL. Ten microliters of each cell suspension were spotted onto YPD agar containing the following stress-inducing agents: 15 mM DTT, 2 µg/mL tunicamycin (Wako Pure Chemical Corporation), 50 µg/mL Calcofluor white (MP Biomedicals), or 250 µg/mL Congo red (Wako Pure Chemical Corporation), and the plates were incubated at 30°C for 2 days. For heat stress, cells were spotted onto YPD agar and incubated at 43°C.

### Detection of spliced and unspliced HAC1 by RT-PCR

*C. auris* cells were grown overnight at 30°C and 1,000 rpm in 10 mL of YPD medium using an RTS-8 (Biosan Ltd., Latvia) until reaching the logarithmic growth phase. Cells were collected by centrifugation and incubated at 30°C for 3 h under the following ER stress conditions: no stress (control), 15 mM DTT, or 2 µg/mL tunicamycin. Total RNA was extracted using ISOGEN (NIPPON GENE) according to the manufacturer’s instructions and further purified using an RNeasy Mini Kit (QIAGEN) with on-column DNase I treatment. cDNA was synthesized from the RNA using ReverTra Ace qPCR RT Master Mix (TOYOBO). RT-PCR was carried out with KOD One PCR Master Mix (TOYOBO) using primers and 50 ng of cDNA as a template. The RT-PCR products were analyzed by electrophoresis on a 1% agarose gel.

### Murine systemic infection model

To analyze the contribution of *HAC1* to virulence, a murine systemic infection model was established according to previously reported methods^26,27^. Briefly, BALB/c mice were immunosuppressed by intraperitoneal injection of cyclophosphamide (Shionogi Inc.) at 200 mg/kg on days -1 and 3 post-infection. A 200-µL suspension of the *C. auris* wild-type, *HAC1* mutant, or *HAC1* complement strain (1.0 ×□10^7^ cells/mouse) was injected intravenously to the immunosuppressed mice. To determine fungal burdens, mice were sacrificed on day 5 post-infection, and the kidneys, livers, and spleens were harvested, homogenized in PBS and plated onto YPD agar plates. Colony-forming units (CFU) were counted after incubation and expressed as CFU per gram of tissue. Each experiment was performed in triplicate.

### Statistical analysis

Statistical analyses were performed using GraphPad Prism, version 10.2.3 (GraphPad Software, San Diego, CA, USA). Survival differences were evaluated using the log-rank (Mantel–Cox) test. For *in vivo* CFU data, the Kruskal–Wallis test followed by Dunn’s multiple comparison test was applied.

## Results

### *In vivo* transcriptome analysis identifies *HAC1* as a virulence-associated gene

To identify genes potentially involved in virulence during *Candida auris* infection, we compared the transcriptomes of cells recovered from an immunosuppressed mouse gastrointestinal infection model with those of cells cultured *in vitro* (Table S1). The intestinal contents and kidneys were collected at 14 days post-infection and defined as *in vivo* (colonization) and *in vivo* (dissemination) samples, respectively. As an *in vitro* control, cells were cultured in yeast nitrogen base medium supplemented with 2% glucose. RNA-seq generated an average of 46.9 million paired-end reads per sample. Mapping rates to the *C. auris* reference genome were high for *in vitro* samples (89–93%) and intestinal samples (71–82%), whereas kidney samples showed markedly lower mapping rates (∼1.6–1.8%), consistent with the predominance of host-derived RNA in disseminated infection tissues.

Principal component analysis (PCA) revealed clear separation between the *in vitro* and *in vivo* samples (Fig. 1a). The first principal component (PC1), which accounted for 83.5% of the total variance, primarily separated *in vitro* samples from host-derived samples. In contrast, colonization and dissemination samples clustered closer together along PC1 but were separated along PC2, suggesting both shared and site-specific transcriptional responses during infection. Replicates from each condition showed broadly consistent clustering patterns. Hierarchical clustering analysis further revealed distinct transcriptional profiles between *in vitro* and *in vivo* conditions (Fig. 1b), with numerous genes upregulated in host-derived samples relative to *in vitro* samples, indicating that the host environment induces substantial transcriptional changes in *C. auris*.

**Figure 1.**
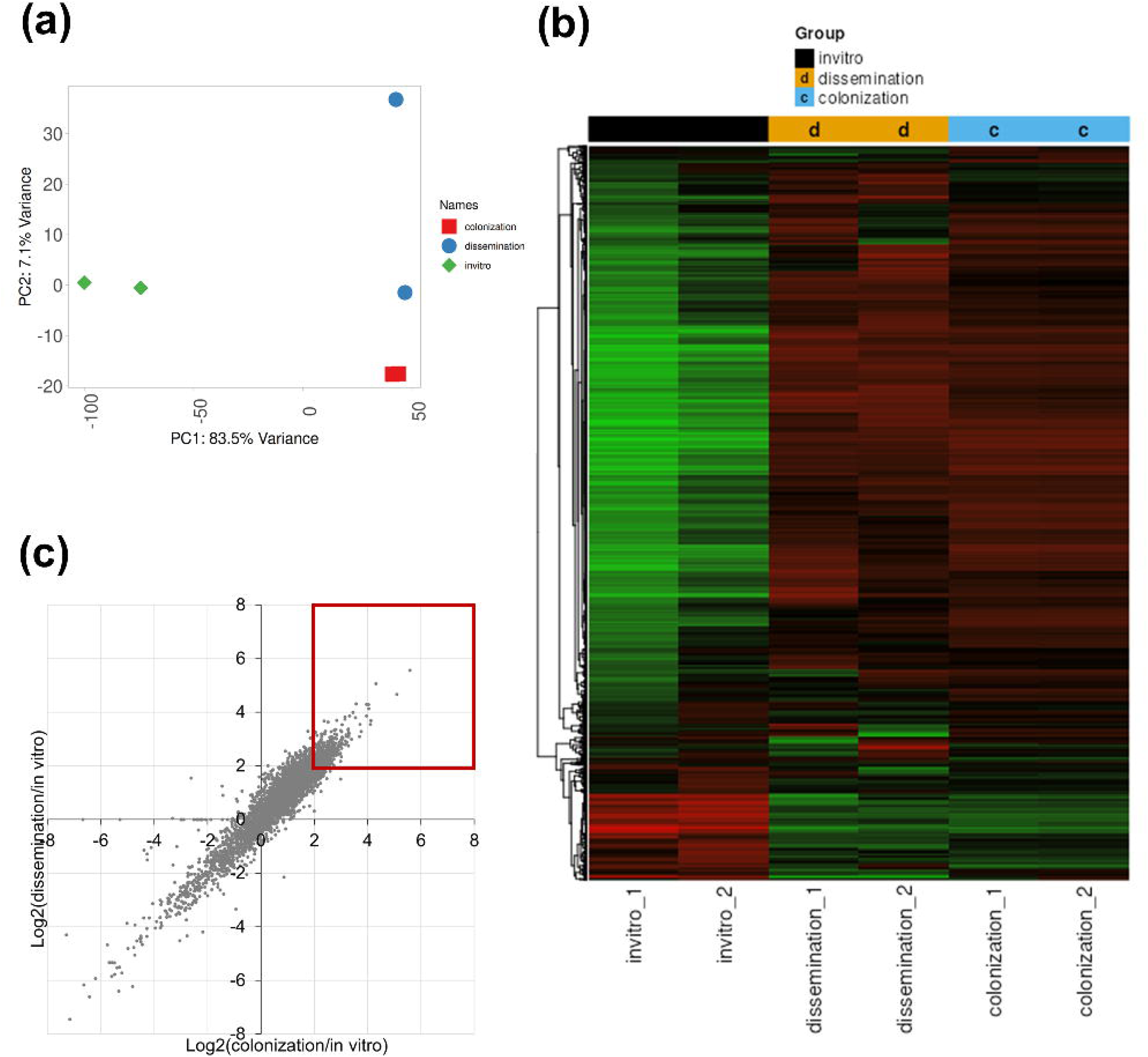
RNA-seq analysis of *in vivo* and *in vitro* samples. (**a**) Principal component analysis of RNA-seq samples obtained from *C. auris* cells recovered from an immunosuppressed mouse infection model and from *in vitro* culture. Green, *in vitro*; red, *in vivo* (colonization); and blue, *in vivo* (dissemination). (**b**) Hierarchical clustering heatmap. Columns represent biological replicates of the *in vitro*, *in vivo* (colonization), and *in vivo* (dissemination) samples. Rows represent genes detected in the RNA-seq dataset. Gene expression levels are shown as relative expression values, with red indicating higher expression and green indicating lower expression. (**c**) Scatter plot comparing gene expression changes in colonization versus dissemination samples relative to *in vitro* samples. Each dot represents a gene. Genes located in the upper-right quadrant (red box) are upregulated under both *in vivo* conditions and were considered candidate virulence-associated genes.

To identify genes potentially associated with infection, we focused on genes that were upregulated in both colonization and dissemination samples relative to the *in vitro* condition. RNA-seq analysis detected expression of 5,594 genes in total. After excluding genes with low baseline expression (FPKM <10 under *in vitro* conditions), 3,846 genes remained for further analysis. Among these, 459 genes showed >4-fold higher expression in colonization samples relative to the *in vitro* condition, while 530 genes showed >4-fold higher expression in dissemination samples. Notably, 338 genes were commonly upregulated under both *in vivo* conditions, suggesting substantial overlap in transcriptional responses during infection (Fig. 1c). From this set, an initial group of 10 genes (B9J08_00015, B9J08_00296, B9J08_01167, B9J08_02474, B9J08_02756, B9J08_03097, B9J08_04418, B9J08_04926, B9J08_05085, and B9J08_05189) was prioritized for deletion screening based on the magnitude of *in vivo* induction and predicted roles in transcriptional regulation, signaling, stress adaptation, and membrane or metabolic functions (Table 2).

### Deletion screening reveals *HAC1* contributes to virulence in *Galleria mellonella*

To determine whether these candidate genes upregulated during infection contribute to virulence, deletion mutants were generated using a CRISPR-Cas9 system. The mutants and the wild-type strain were injected into *G. mellonella* larvae, and survival was monitored for 7 days (Fig. 2). Larvae infected with the wild-type strain showed decreased survival beginning on day 1 post-infection, with a 7-day survival rate of 10%. In contrast, larvae infected with the B9J08_00296 deletion mutant exhibited delayed mortality and a significantly higher 7-day survival rate of 50%. The median survival time was 2 days for the wild-type group and 7 days for the B9J08_00296 deletion mutant group. Infection with the B9J08_00296-complemented strain restored virulence to levels comparable to those of the wild-type strain (Fig. S1), confirming that the observed phenotype was attributable to B9J08_00296 deletion. Although deletion mutants of B9J08_03097 and B9J08_05085 showed statistically significant differences compared with the wild-type strain, their effects on survival were relatively modest. The other mutants did not exhibit significant differences. According to database information, B9J08_00296 encodes Hac1, a transcription factor that regulate the UPR pathway. Together, these results indicate that, among the candidates examined, Hac1 plays a significant role in virulence in the *G. mellonella* infection model.

**Figure 2.**
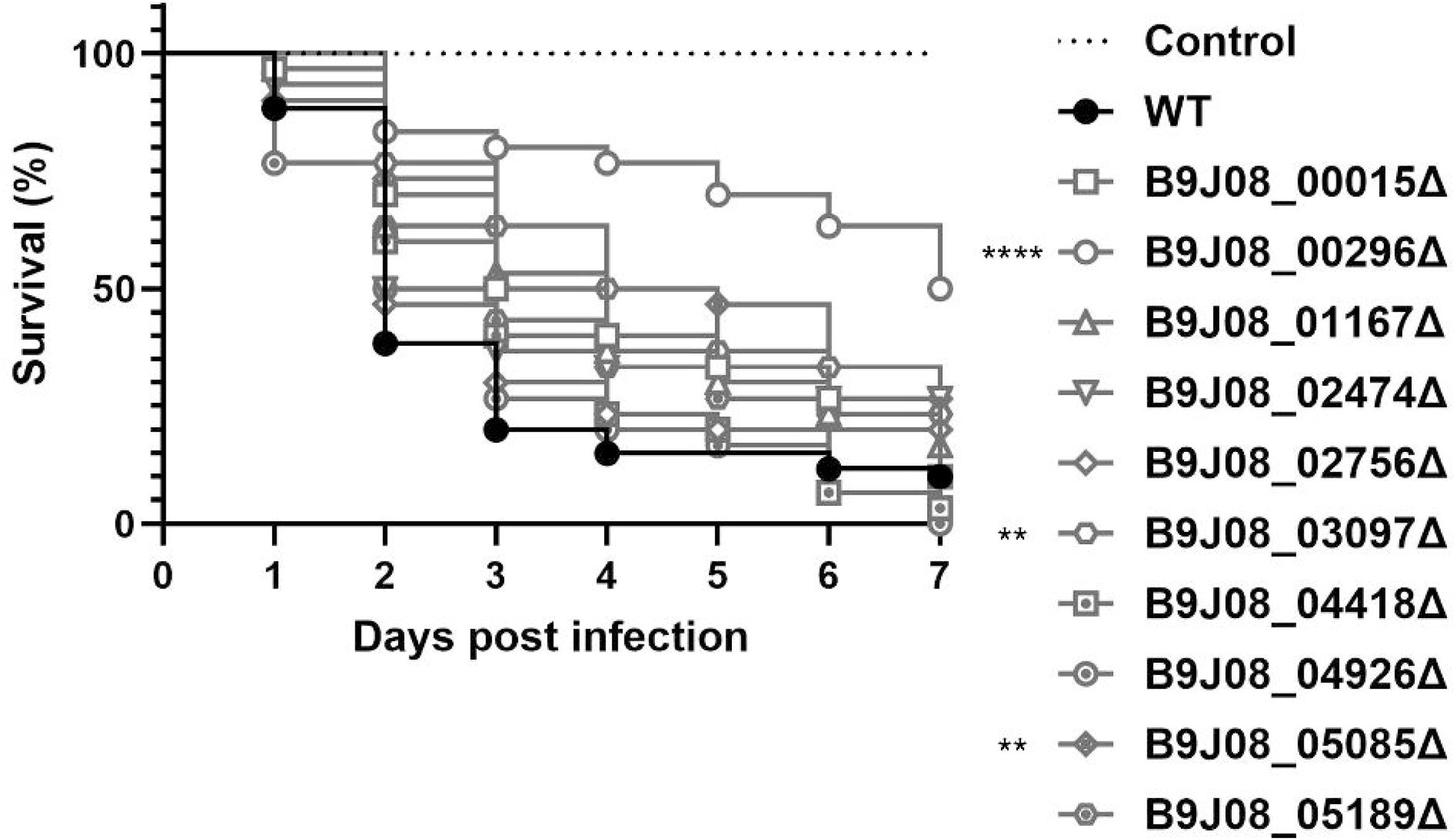
Deletion screening of candidate genes in the *Galleria mellonella* infection model. Larvae of *G. mellonella* were infected with the wild-type (WT) strain or gene deletion mutants (B9J08_00015Δ, B9J08_00296Δ, B9J08_01167Δ, B9J08_02474Δ, B9J08_02756Δ, B9J08_03097Δ, B9J08_04418Δ, B9J08_04926Δ, B9J08_05085Δ, and B9J08_05189Δ) and monitored daily for 7 days post infection. Survival differences were evaluated using the log-rank (Mantel–Cox) test. **** *P*□<□0.0001;; ** *P*□<□0.01.

### ER stress induces unconventional splicing of *HAC1* mRNA

To investigate the molecular mechanism of *HAC1* activation, we first investigated the susceptibility of *C. auris* to ER stress. Growth assays on YPD agar containing tunicamycin, an inhibitor of N-linked glycosylation, or dithiothreitol (DTT), which disrupts disulfide bond formation, were performed to evaluate stress susceptibility among the clade strains (Fig. 3a). Under DTT treatment, AR 0390 (clade I) and AR 0385 (clade IV) showed relatively low sensitivity, whereas AR 0381 (clade II) and AR 1097 (clade V) were more sensitive, with AR 0384 (clade III) displaying intermediate sensitivity. Under tunicamycin treatment, AR 0385 (clade IV) exhibited the lowest sensitivity, followed by AR 1097 (clade V), whereas AR 0390 (clade I) and AR 0384 (clade III) showed moderate sensitivity, and AR 0381 (clade II) was the most sensitive.

**Figure 3.**
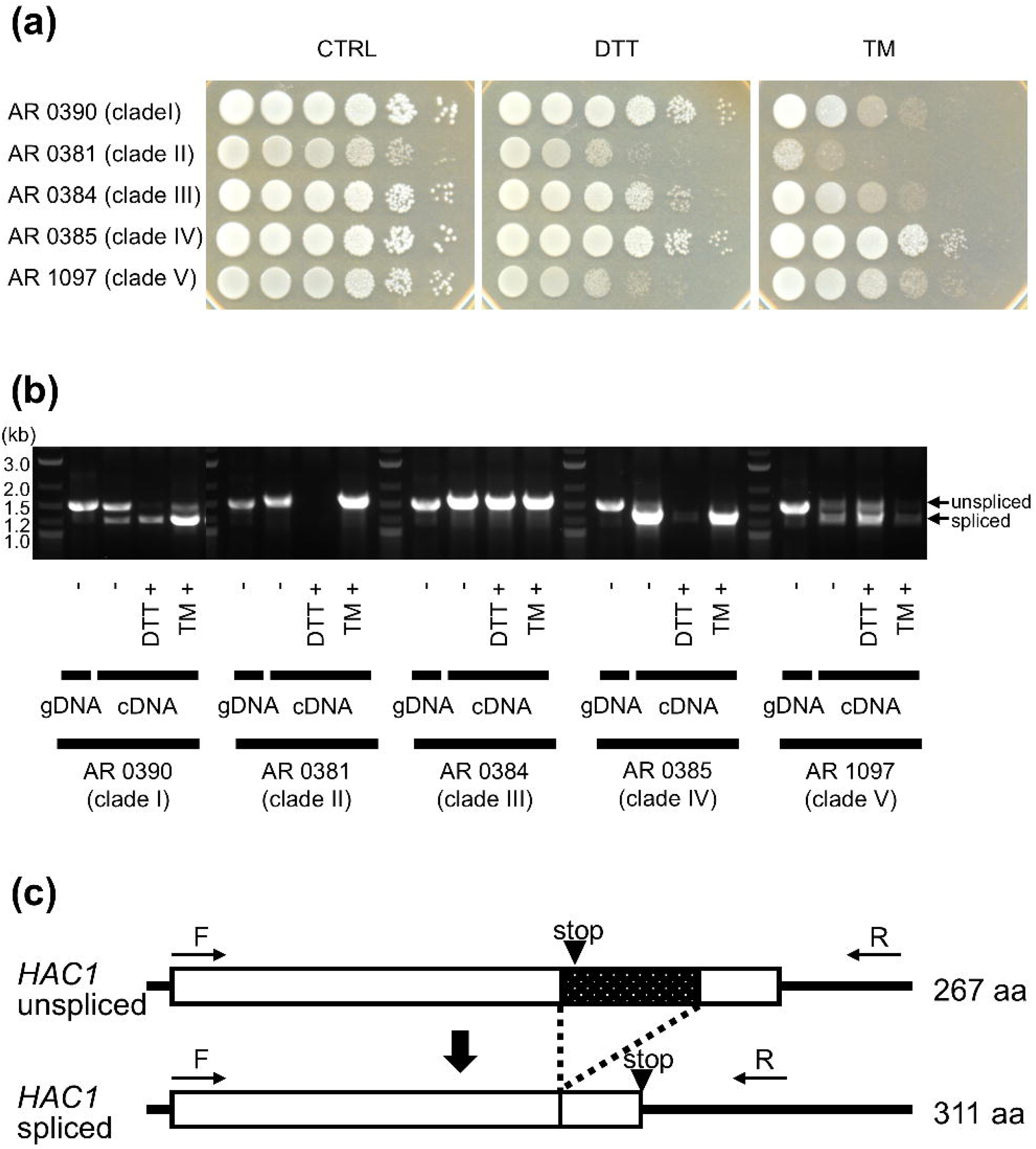
Unconventional splicing of *HAC1* mRNA under ER stress conditions in *Candida auris*. (**a**) Growth assay of *C. auris* clade strains under ER stress conditions. Serial 10-fold dilutions of cells (1.0 ×□10^6^ to 1.0 ×□10 cells/spot) from each clade strain (AR 0390, AR 0381, AR 0384, AR 0385, and AR 1097) were spotted onto YPD agar plates containing 15 mM DTT or 2 µg/mL tunicamycin and incubated at the indicated temperature. (**b**) RT-PCR analysis of *HAC1* mRNA splicing in *C. auris*. Cells were grown under unstressed conditions or in the presence of ER stress–inducing agents. RT-PCR products corresponding to unspliced (larger) and spliced (smaller) *HAC1* transcripts were detected by agarose gel electrophoresis. -, No stress; DTT, dithiothreitol; and TM, tunicamycin. (**c**) Schematic representation of the *C. auris HAC1* mRNA splicing event. RT-PCR and sequencing analyses identified a 287-bp region that is removed during splicing (dotted region).

Then, we investigated whether *HAC1* mRNA undergoes unconventional splicing to generate the active Hac1 transcript in *C. auris* by RT-PCR analysis using clade I-V strains grown under unstressed or ER stress conditions (Fig. 3b). Primers were designed to amplify a region spanning the predicted intron based on the annotated *HAC1* sequence (Table 1). In all strains, PCR using genomic DNA as a template yielded an approximately 1.5 kb band, corresponding to the unspliced form. In contrast, RT-PCR using cDNA from unstressed cells produced two distinct bands: an approximately 1.5 kb band corresponding to the unspliced form and a smaller band of approximately 1.2 kb corresponding to the spliced form. In AR 0390 (clade I), a fraction of *HAC1* mRNA was constitutively spliced even under basal conditions, and ER stress induced by tunicamycin or DTT increased the abundance of the spliced form. In AR 0381 (clade II), no clear spliced band was detected under unstressed conditions, and tunicamycin treatment produced only a faint smear-like signal. Under DTT treatment, no *HAC1* amplification product was observed. In AR 0384 (clade III), the spliced form was barely detectable under all conditions. In contrast, AR 0385 (clade IV) exhibited an intense spliced band even under unstressed conditions, and this predominance became more evident after tunicamycin or DTT treatment. In AR 1097 (clade V), both unspliced and spliced bands were detected at comparable levels under unstressed and ER stress conditions.

**Table 1.**
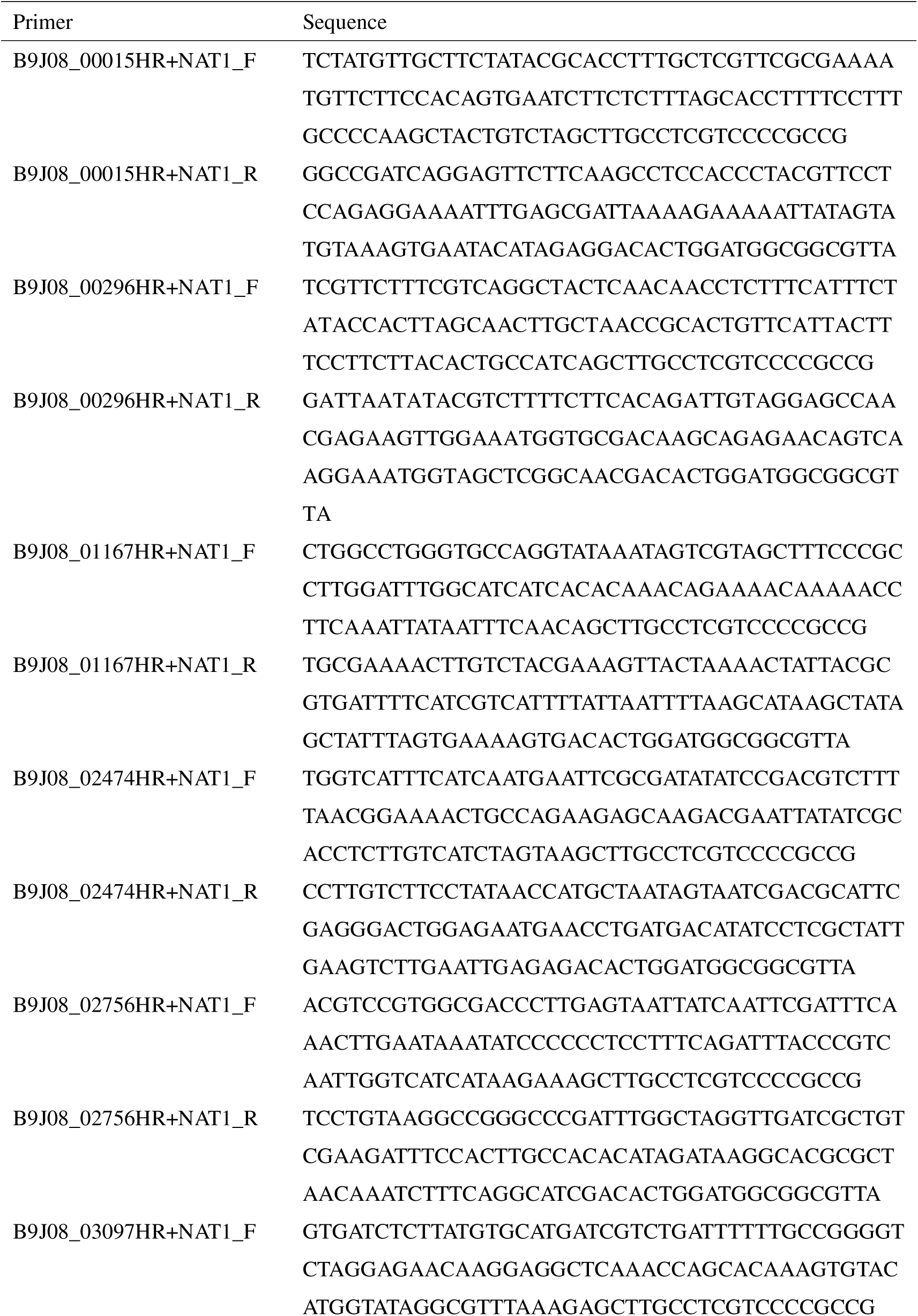
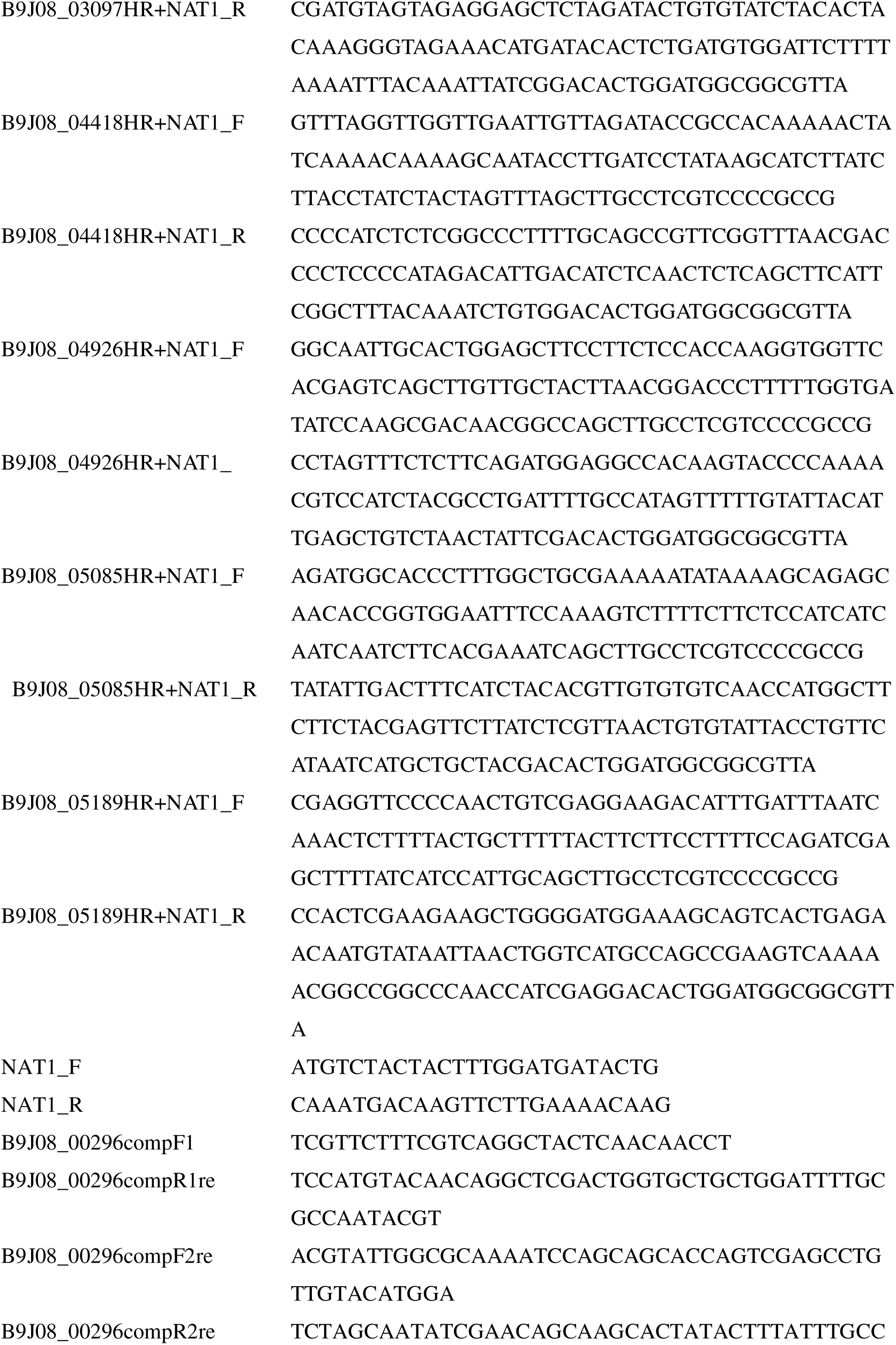
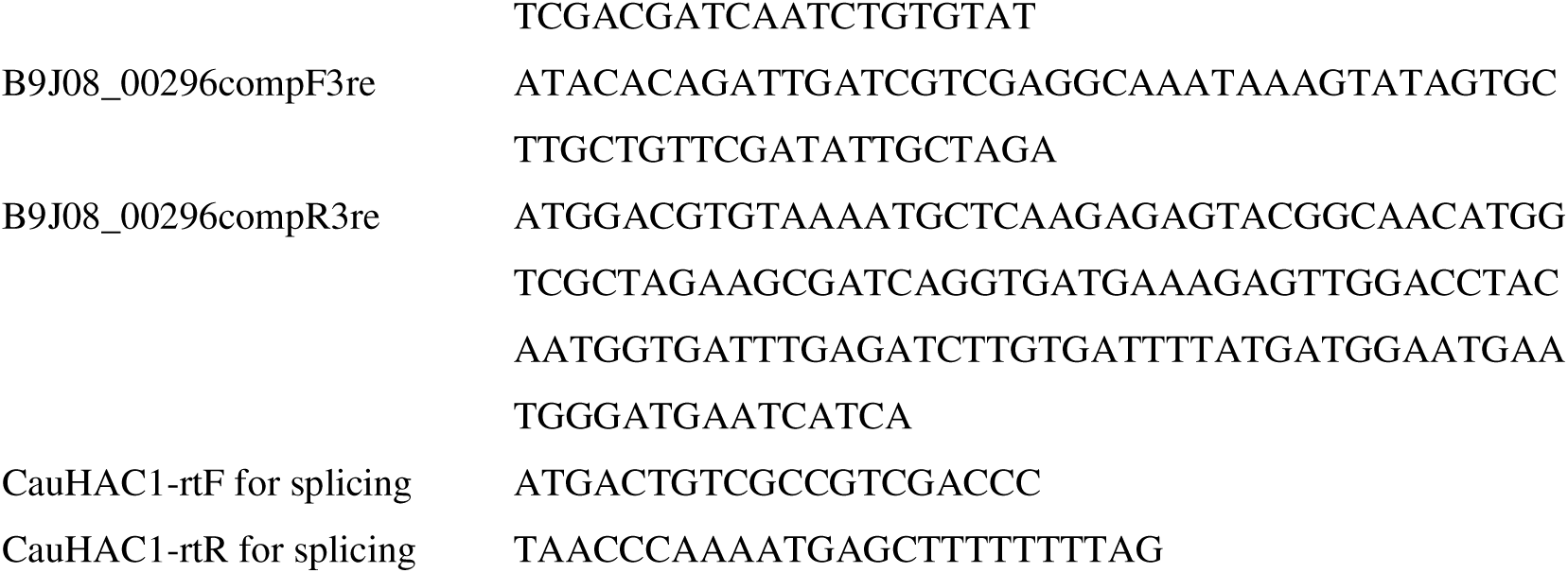
Sequences of primers.

**Table 2.** Candidate genes upregulated during infection.

| Gene ID | FPKM |  |  | Log<sub>2</sub>Fold change |  | Candida Genome Database annotation |
| --- | --- | --- | --- | --- | --- | --- |
|  | In vitro | Colonization | Dissemination | Colonization<br>/ in vitro | Dissemination<br>/ in vitro |  |
| B9J08_00015 | 11.66 | 104.05 | 88.52 | 3.16 | 2.92 | Ortholog(s) have role in GPI anchor biosynthetic process, intracellular manganese ion homeostasis and endoplasmic reticulum, fungal-type vacuole membrane localization |
| B9J08_00296 | 101.50 | 1329.75 | 1185.94 | 3.71 | 3.55 | Ortholog(s) have DNA-binding transcription factor activity, RNA polymerase II-specific activity |
| B9J08_01167 | 2342.05 | 19309.93 | 19681.59 | 3.04 | 3.07 | Has domain(s) with predicted NAD binding, NADP binding, oxidoreductase activity, acting on the aldehyde or oxo group of donors, NAD or NADP as acceptor activity and role in glucose metabolic process |
| B9J08_02474 | 14.98 | 83.84 | 84.18 | 2.48 | 2.49 | Ortholog(s) have phospholipase A2 activity, role in cardiolipin acyl-chain remodeling, cardiolipin metabolic process and mitochondrial inner membrane, mitochondrion localization |
| B9J08_02756 | 89.05 | 385.71 | 404.18 | 2.11 | 2.18 | Ortholog of *C. albicans* SC5314 : C7_00360W_A/DFI1, *C. dubliniensis* CD36 : Cd36_70570, *C. parapsilosis* CDC317 : CPAR2_301600 and *Debaryomyces hansenii* CBS767 : DEHA2F13376g |
| B9J08_03097 | 25.61 | 169.61 | 121.84 | 2.73 | 2.25 | Ortholog(s) have DNA-binding transcription repressor activity, RNA polymerase II-specific, histone H3K36 demethylase activity, histone H3K9 demethylase activity, sequence-specific DNA binding activity |
| B9J08_04418 | 11.96 | 572.24 | 563.14 | 5.58 | 5.56 | Ortholog(s) have protein kinase activity, protein serine/threonine kinase activity |
| B9J08_04926 | 24.32 | 196.90 | 169.62 | 3.02 | 2.80 | Has domain(s) with predicted triglyceride lipase activity and role in lipid catabolic process |
| B9J08_05085 | 1502.32 | 14570.71 | 15526.91 | 3.28 | 3.37 | Has domain(s) with predicted oxidoreductase activity, zinc ion binding activity |
| B9J08_05189 | 10.10 | 200.19 | 338.76 | 4.31 | 5.07 | Ortholog(s) have role in intracellular signal transduction |

Sequencing of the RT-PCR products revealed that the spliced region comprised 287 bp, spanning from 31-bp upstream to 256-bp downstream of the annotated stop codon in the Candida Genome Database (Fig. 3b). These findings suggest that the actual translation stop site may differ from the annotated ORF, potentially producing a 311-amino-acid Hac1 protein distinct from the currently predicted sequence. Collectively, these observations suggest a potential relationship between *HAC1* splicing levels and ER stress sensitivity among *C. auris* clades.

### *HAC1* contributes to ER stress and heat stress tolerance

To investigate the functional role of *HAC1*, growth assays were performed using the wild-type strain, the *HAC1* deletion mutant, and the complemented strain under various stress conditions, including ER stress (tunicamycin and DTT), cell wall stress (Calcofluor white and Congo red), and heat stress (Fig. 4). The *HAC1* deletion mutant strain showed reduced growth on YPD agar containing tunicamycin compared with the wild-type strain. While no growth difference was observed at 30°C, the *HAC1* deletion mutant showed impaired growth at elevated temperature (43°C). In all cases, the complemented strain restored growth to wild-type levels. In contrast, no increased sensitivity was observed in the *HAC1* deletion mutant in the presence of DTT, Calcofluor white, or Congo red. These results suggest that *HAC1* plays a specific role in adaptation to ER stress and heat stress, but not to cell wall stress, in *C. auris*.

**Figure 4.**
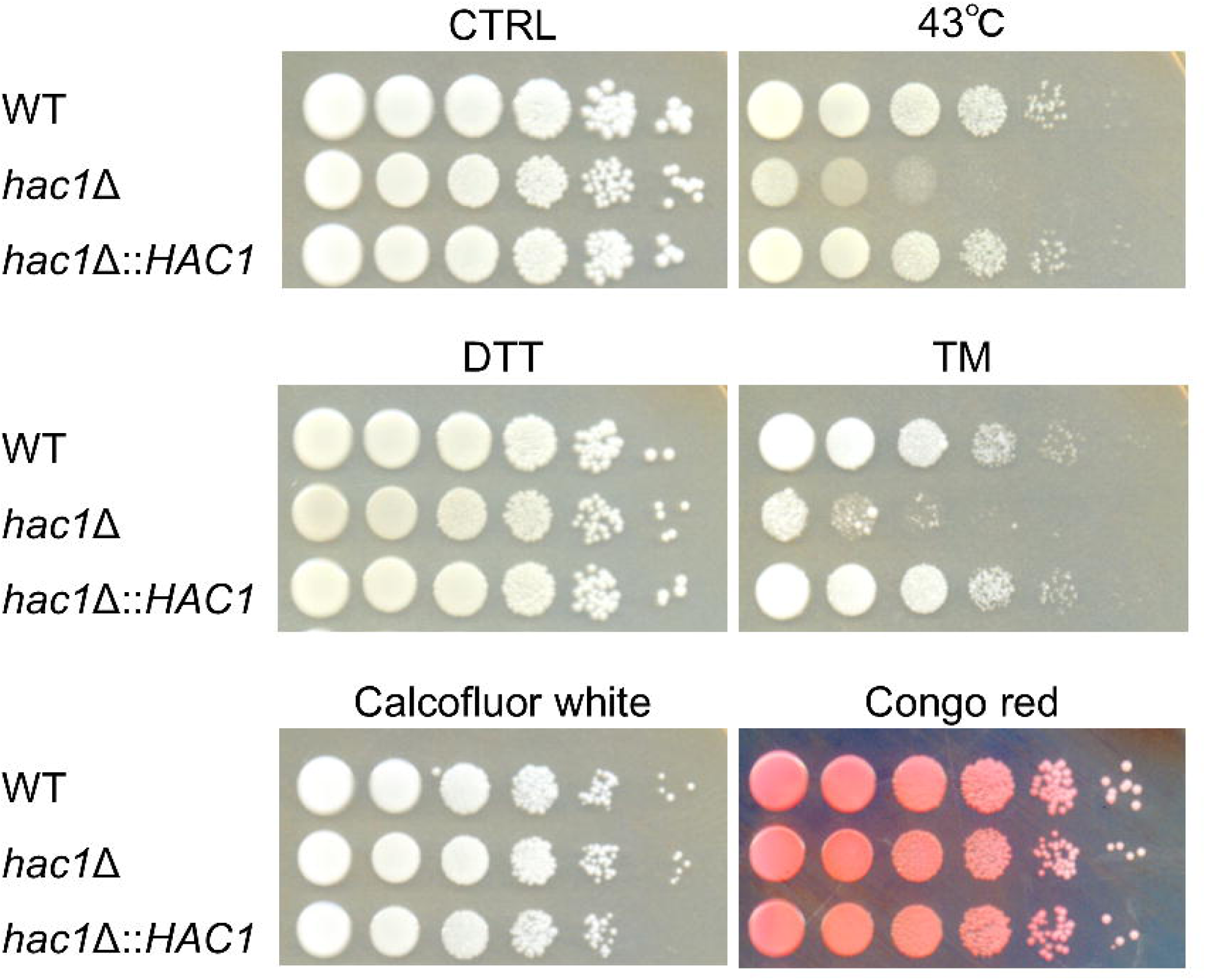
Sensitivity of *C. auris HAC1* deletion mutant to various stresses. Growth assay of the wild-type (WT), *HAC1* deletion mutant (*hac1*Δ), and complemented strain (*hac1*Δ::*HAC1*) under the indicated conditions: CTRL, no stress; 43°C, YPD agar incubated at 43°C; DTT, 15 mM dithiothreitol; TM, 2 µg/mL tunicamycin; Calcofluor white, 50 µg/mL Calcofluor white; Congo red, 250 µg/mL Congo red.

### *HAC1* contributes to virulence in an immunosuppressed mouse model

To further evaluate the role of *HAC1* in mammalian virulence, immunosuppressed mice were intravenously infected with the wild-type strain, the *HAC1* deletion mutant, or the complemented strain, and fungal burdens in the kidneys, liver, and spleen were quantified on day 5 post-infection (Fig. 5). In the kidneys, the fungal burden of the wild-type strain was 7.44 ± 0.41 log CFU/g, whereas that of the *HAC1* deletion mutant was reduced by more than one order of magnitude (5.65 ± 0.47 log CFU/g), with statistical significance. The complemented strain showed a fungal burden of 7.08 ± 0.52 log CFU/g. In the liver, the wild-type strain yielded 4.58 ± 0.57 log CFU/g, compared with 2.82 ± 0.57 log CFU/g (mutant) for the *HAC1* deletion mutant and 4.29 ± 0.94 log CFU/g for the complemented strain. A similar trend was observed in the spleen, where fungal burdens were 6.23 ± 0.20 log CFU/g, 5.26 ± 0.12 log CFU/g, and 6.15 ± 0.15 log CFU/g for the wild-type, mutant, and complemented strains, respectively. Together, these results indicate that *HAC1* contributes to systemic virulence in *C. auris*.

**Figure 5.**
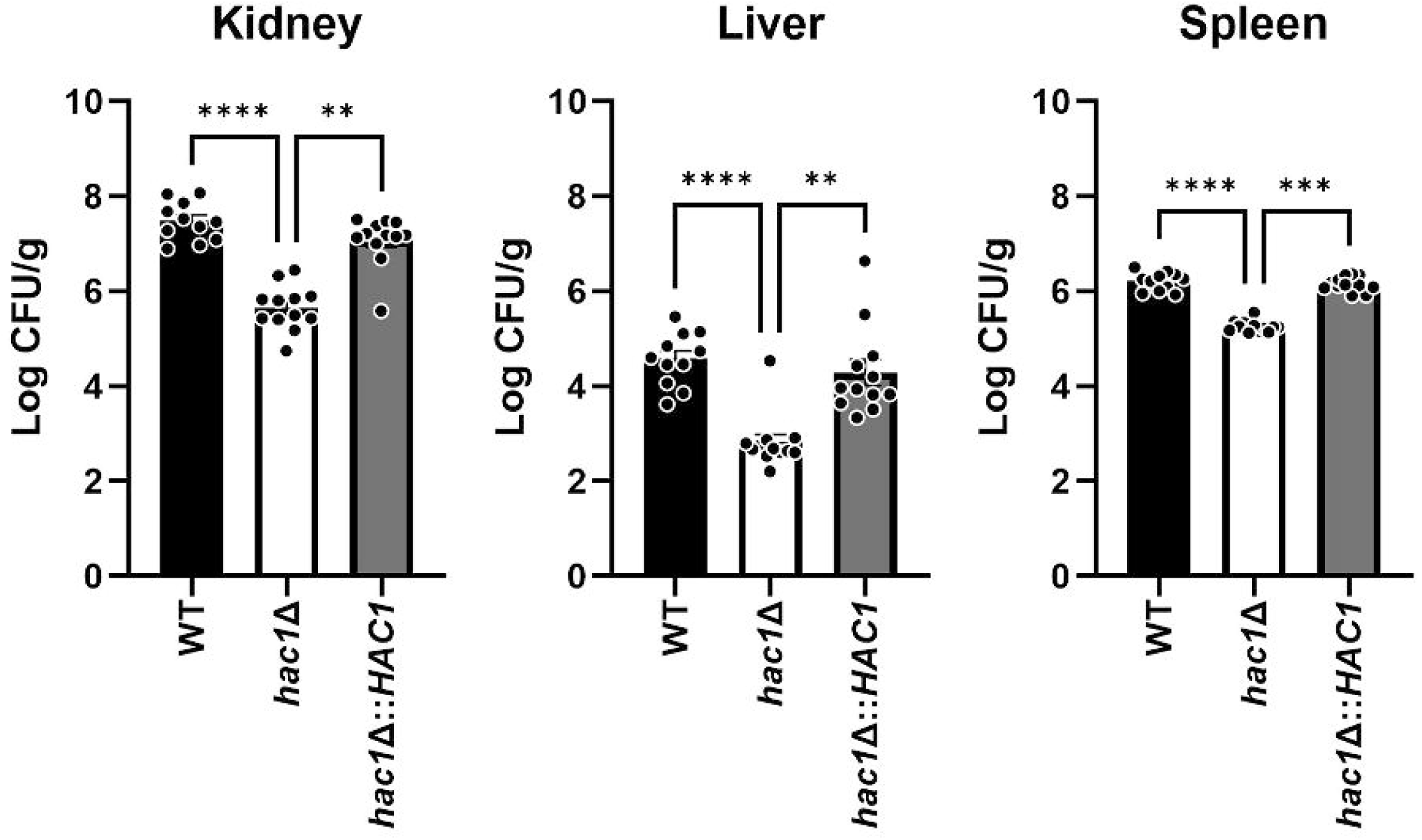
Fungal burdens of *C. auris HAC1* deletion mutant in the immunocompromised mouse infection model. Fungal burdens were determined by plating homogenates of kidney, liver, and spleen tissues and are presented as log CFU per gram of tissue. Black, the wild-type strain (WT); white, *HAC1* deletion mutant (*hac1*Δ); or gray, complemented strain (*hac1*Δ::*HAC1*). Each dot represents an individual mouse, and bars indicate the mean ± standard error of the mean. **** *P*□<□0.0001; *** *P*□<□0.001; ** *P*□<□0.01.

## Discussion

In this study, *in vivo* transcriptome analysis identified *HAC1* as a virulence-associated gene in *C. auris*. Examination of genes commonly induced during both colonization and dissemination enabled us to identify genes that were consistently induced across distinct infection stages and therefore more likely to represent core determinants of host adaptation rather than niche-specific responses. Among these genes, *HAC1* was particularly notable because of its established role in the UPR.

Analysis of *HAC1* mRNA splicing in *C. auris* revealed an unconventional splicing event. Although the Candida Genome Database does not annotate an intron in *HAC1*, RT-PCR and sequencing analyses identified a 287-bp unconventional splicing that is removed under ER stress conditions. Because this excised region spans the currently annotated ORF boundary, the translated region of *C. auris HAC1* may require re-evaluation. Interestingly, a proportion of *HAC1* mRNA appeared to be spliced even under non-stress conditions, suggesting that basal *HAC1* activation may occur constitutively in *C. auris*. The increase in the spliced form following tunicamycin or DTT treatment further indicates that *HAC1* splicing is enhanced under ER stress conditions. These observations suggest that stress-responsive HAC1 activation is retained in *C. auris*, although the basal level of splicing activity may be relatively high. In addition, differences in splicing dynamics were also observed among clades. Clade I and IV isolates showed relatively strong spliced bands in the presence or absence of ER stress, whereas clade II and III isolates exhibited more limited splicing induction. These findings suggest that basal UPR activity or its regulation may differ across *C. auris* clades.

Functional analyses further supported an important role for *HAC1*. The *HAC1* deletion mutant showed increased sensitivity to tunicamycin and elevated temperature, whereas no clear differences were observed in response to cell wall stressors such as Calcofluor white and Congo red. In both *G. mellonella* and immunosuppressed mouse infection models, loss of *HAC1* resulted in reduced virulence, as indicated by prolonged survival of infected larvae and decreased fungal burdens in mice. Notably, the *HAC1* deletion mutant also exhibited marked growth impairment under high-temperature conditions, further indicating a role for *HAC1* in thermotolerance. This thermotolerance defect may be particularly relevant during infection, because mammalian hosts impose a thermal environment that is substantially higher than typical environmental conditions. Increased temperature can disrupt protein folding in the ER and increase the burden on ER quality-control systems^28^. In fungal pathogens, the UPR and other proteostasis mechanisms are required to maintain protein homeostasis during thermal stress and host temperature adaptation^29^. Together, these findings support a critical role for *HAC1* in ER stress adaptation, temperature tolerance, and survival within the host environment.

Comparison with other *Candida* species highlights both the conservation and diversification of the UPR pathway. In *C. albicans*, *HAC1* mRNA undergoes unconventional splicing mediated by the ER-resident sensor Ire1, and Hac1 has been shown to contribute to ER stress adaptation as well as morphogenesis and virulence-related traits^20^. In contrast, *Candida glabrata* appears to have diverged from this canonical organization: although Ire1 is required for ER stress adaptation, the stress response occurs largely independently of Hac1, suggesting substantial rewiring of the classical Ire1–Hac1 pathway^30^. *Candida parapsilosis* provides another variation, as it possesses an unusually long *HAC1* intron, and deletion of *HAC1* in this species increases sensitivity to ER stress, cell wall stress, and certain antifungal agents^31^. Thus, even within the genus *Candida*, both the mechanism and physiological consequences of *HAC1* splicing vary substantially. The present findings indicate that *C. auris* also exhibits inducible *HAC1* mRNA splicing, suggesting that a splicing-based mechanism of UPR activation is retained in this species. Functionally, the relatively restricted stress phenotypes observed in the *HAC1* deletion mutant further suggest that the downstream output of the UPR pathway may differ from that in other *Candida* species. In particular, the absence of clear sensitivity to DTT or cell wall stressors in the mutant contrasts with observations in *C. albicans* and *C. parapsilosis*, where disruption of the UPR has broader effects on ER stress tolerance and virulence-associated processes. These differences may reflect species-specific rewiring of stress response networks downstream of *HAC1*.

Several limitations should be noted. First, the upstream regulatory mechanism responsible for *HAC1* splicing was not directly examined in this study, and it therefore remains unclear whether *HAC1* activation in *C. auris* is strictly dependent on Ire1. Second, the splicing analysis was based largely on semi-quantitative RT-PCR, and more precise quantitative approaches would help to strengthen these observations. Third, both the RNA-seq analysis and clade comparison were performed using a limited number of strains, and analysis of a broader strain set will be necessary to evaluate intraspecies diversity.

In conclusion, this study identifies *HAC1* as a virulence-associated gene in *C. auris* and supports a role for the UPR pathway in adaptation and survival in the host environment. The observed clade-dependent differences in splicing dynamics further suggest diversity in stress response regulation within *C. auris*. Understanding the evolutionary rewiring of the UPR pathway across *Candida* species may provide a useful framework for developing antifungal strategies targeting ER stress adaptation.

## Acknowledgments

We are grateful to Dr. Hiroji Chibana and Michiyo Okamoto (Chiba University) for kindly providing the plasmid originally described by Sherin Shahana et al. We also appreciate the Centers for Disease Control and Prevention for supplying the Antibiotic Resistance Isolate Bank *C. auris* isolates in this study.

## Data Repository

The RNA-seq data have been deposited in the DDBJ Sequence Read Archive under BioProject accession number PRJDB40537.

## Author contributions

Sayoko Oiki (Conceptualization, Data curation, Formal analysis, Funding acquisition, Investigation, Methodology, Validation, Visualization, Writing—original draft), Masahiro Abe (Conceptualization, Formal analysis, Funding acquisition, Investigation, Methodology, Writing—review & editing), Ai Hirasawa (Investigation), Ami Koizumi (Formal analysis), Amato Otani (Formal analysis), Takayuki Shinohara (Formal analysis), and Yoshitsugu Miyazaki (Conceptualization, Funding acquisition, Project administration, Resources, Supervision, Writing—review & editing).

