## Supplementary material for "*HAC1* contributes to stress adaptation and virulence in the emerging fungal pathogen *Candida auris*": Fig. S1

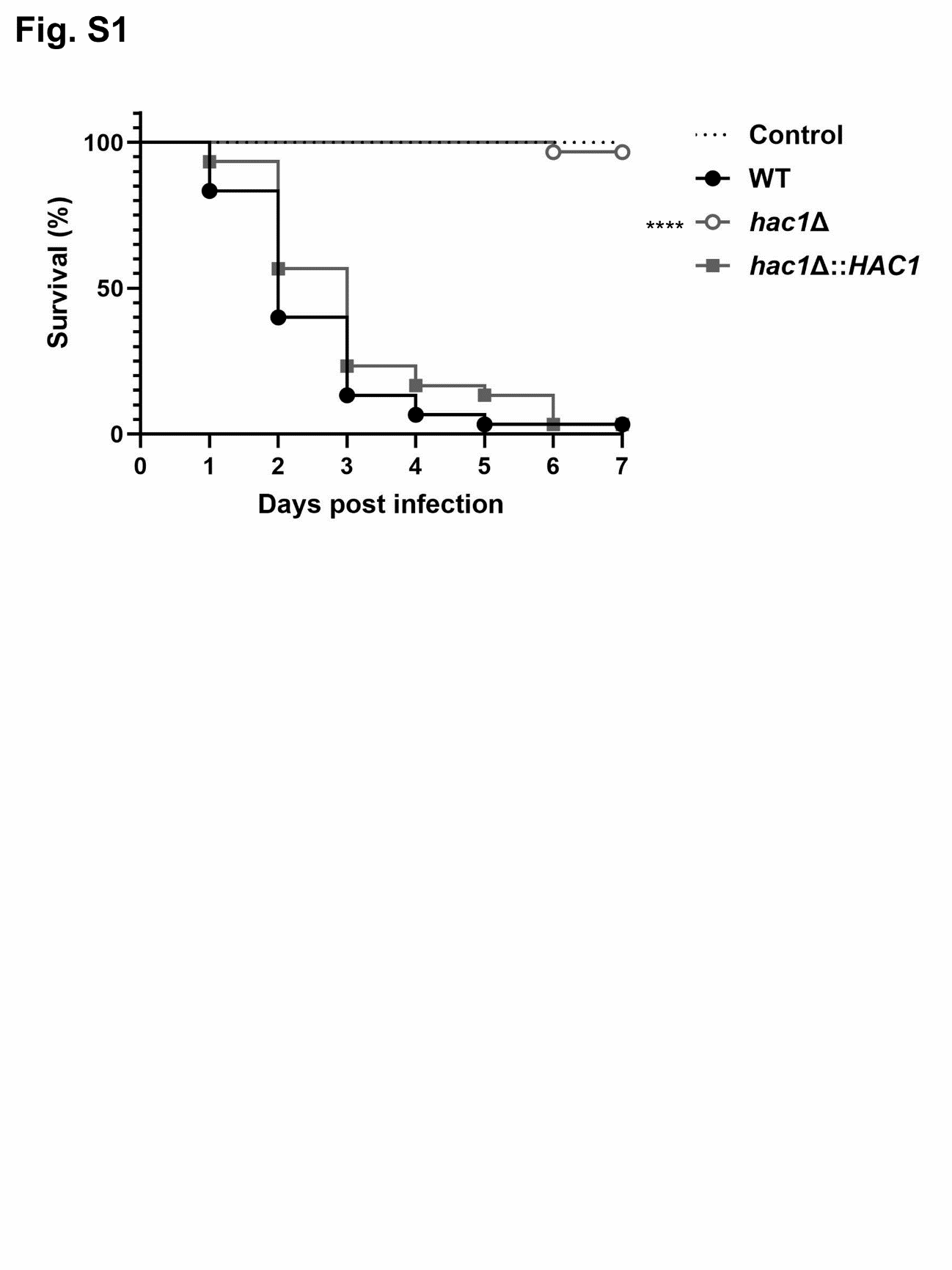

**Figure S1 Survival of *G. mellonella* larvae infected with *C. auris*.** Black, AR 0390 wild-type strain (WT); white, HAC1 deletion mutant (HAC1Δ); gray, HAC1 complemented strain (HAC1cp), dotted line; control.
